# Outer membrane-deprived cyanobacteria liberate periplasmic and thylakoid luminal components that support the growth of heterotrophs

**DOI:** 10.1101/2020.03.24.006684

**Authors:** Seiji Kojima, Yasuaki Okumura

## Abstract

Chloroplasts originate from endosymbiosis of a cyanobacterium within a heterotrophic host cell. Establishing endosymbiosis requires the translocation across its envelope of photosynthetic products generated inside the once free-living cyanobacterium to be exploited by host metabolism. However, the nature of this translocation event is unknown. We previously found that most cyanobacterial outer membrane components were eliminated during the primitive stage of chloroplast evolution, suggesting the importance of evolutionary changes of the outer membrane. Here, we removed the outer membrane from *Synechocystis* sp. PCC 6803 by disrupting the physical interaction with peptidoglycan, and characterized the effects on cell function. Outer membrane-deprived cells liberated diverse substances into the environment without significantly compromising photoautotrophic growth. The amount of liberated proteins increased to ~0.35 g/L within five days of culture. Proteomic analysis showed that most liberated proteins were periplasmic and thylakoid luminal components. Connectivity between the thylakoid lumen-extracellular space was confirmed by findings that an exogenous hydrophilic oxidant was reduced by photosynthetic electron transport chain on the thylakoid membrane. Metabolomic analysis detected the release of nucleotide-related metabolites at concentrations around 1 μM. The liberated materials supported the proliferation of heterotrophic bacteria. These findings show that breaching the outer membrane, without any manipulations to the cytoplasmic membrane, converts a cyanobacterium to a chloroplast-like organism that conducts photosynthesis and releases its biogenic materials. This conversion not only represents a potential explanation why the outer membrane markedly changed during the earliest stage of chloroplast evolution, but also provides the opportunity to harness cyanobacterial photosynthesis for biomanufacturing processes.

**SIGNIFICANCE STATEMENT:** Although it is well accepted that chloroplasts stem from endosymbiosis of a cyanobacterium within a heterotrophic host cell, the issue of how photosynthetic products generated inside a formerly free-living cyanobacterium are translocated across its envelope and exploited by host metabolism has been little addressed. Here we show that breaching the cyanobacterial outer membrane barrier converts a cyanobacterium to a chloroplast-like organism that conducts photosynthesis and releases its diverse biogenic materials into its external environment, which sustains the growth of heterotrophic organisms. This conversion represents a possible example of metabolic exploitation of cyanobacterial photosynthesis. Further, this “quasi-chloroplast” provides a potential opportunity for industrial application such as producing feedstock for biomanufacturing processes that harnesses heterotrophic bacteria.

## Introduction

Chloroplasts originate from endosymbiosis of a bacterium belonging to the phylum Cyanobacteria, the Gram-negative bacteria that conduct oxygenic photosynthesis, within a heterotrophic host cell (1–3). The primary endosymbiosis event, estimated to have occurred approximately 2.1 billion years ago, gave rise to the Archaeplastida comprising glaucophytes, red algae (Rhodophyta), green algae, and land plants (Viridiplantae). Through secondary endosymbiosis, in turn, red or green algae were incorporated into other heterotrophic hosts and further spread chloroplasts to haptophytes, cryptophytes, euglenids, and various other organisms. Chloroplasts are now central to the primary production on the earth, sustaining biological ecosystems.

Establishing an endosymbiotic relationship requires the translocation of photosynthetic products generated inside the once free-living cyanobacterium across its envelope to be exploited by the host’s metabolism. Although membrane transport of the chloroplast from extant plants and algae is well studied (4), the nature of the earliest mechanism of translocation during the initial stage of endosymbiosis is far less addressed, despite the wealth of accumulating genetic information. An obvious difficulty is ascribed to the facts that chloroplast genomes underwent a drastic reduction in size compared to that of cyanobacteria, and extant genomes of plants, as well as those of cyanobacteria, have accepted massive horizontal gene transfers. These complex genomic contexts present a difficult challenge for phylogenetics in the efforts to deduce key genetic change(s) relevant to establishing the translocation mechanism (1). It was argued that genes derived from *Chlamydiae* contributed to the generation of an export pathway for photosynthetic products, e.g., hexose phosphate, across the endosymbiont’s inner membrane and thereby enabled metabolic exploitation of the endosymbiont (5, 6). However, this remains an open question (7, 8).

Glaucophytes retain primitive chloroplasts (referred to as muroplasts) with a cyanobacteria-like envelope structure comprising the inner membrane, peptidoglycan, and outer membrane (6, 9, 10). Therefore it provides opportunities to illustrate the initial stage of chloroplast evolution. We previously discovered that the outer membrane of the muroplast predominantly comprises the noncyanobacterial lineage proteins CppS and CppF (11). Coincidentally, the most abundant outer membrane protein of cyanobacteria, i.e., Slr1841 of *Synechocystis* sp. PCC 6803 (hereafter referred to as PCC 6803) and its homologs which represent approximately 80% of all the proteins in cyanobacterial outer membrane preparations (Fig. 1A) (12), are not conserved in an extant plant lineage. This replacement most likely rendered the ancestral cyanobacterial outer membrane highly permeable to diverse organic compounds for reasons as follows: (i) Slr1841-like proteins function as an inorganic ion-permeating channel and that prevents the transport of organic compounds (12), whereas CppS and CppF serve as a nonspecific diffusion channel that allows permeation of compounds with *M_r_* ~1,000 (11)]. (ii) Depriving Slr1841-like proteins, which bind to the peptidoglycan via their periplasm-exposed S-layer homologous (SLH) domains (11, 12), is expected to disrupt the integrity of the outer membrane because Gramnegative bacteria generally requires outer membrane–peptidoglycan linkage to maintain the permeability barrier (13–16). Strikingly, the latter scenario can be deduced by another evolutionary change involving cyanobacterial peptidoglycan-linked polysaccharides, which are required for interaction with the SLH domain (Fig. 1A) but are missing from muroplast peptidoglycan (9). The occurrence of extensive evolutionary changes to the outer membrane is also evident by the fact that only three of 20 outer membrane proteins, identified by a proteomic analysis of PCC 6803 (17), are conserved in the extant plant lineage (Table S1). In contrast, 27 of 57 cytoplasmic membrane proteins of PCC 6803 occur in plants (Table S2) (18). Because the outer membrane functions as the outermost barrier separating the cell interior from the external environment (12, 19, 20), breaching its permeability barrier presumably causes the release of diverse cellular organic compounds, which are originally generated as products of photosynthesis, into the external environment that corresponds to the host cytosol in case the cyanobacterium is in an endosymbiotic situation.

**Fig. 1.**
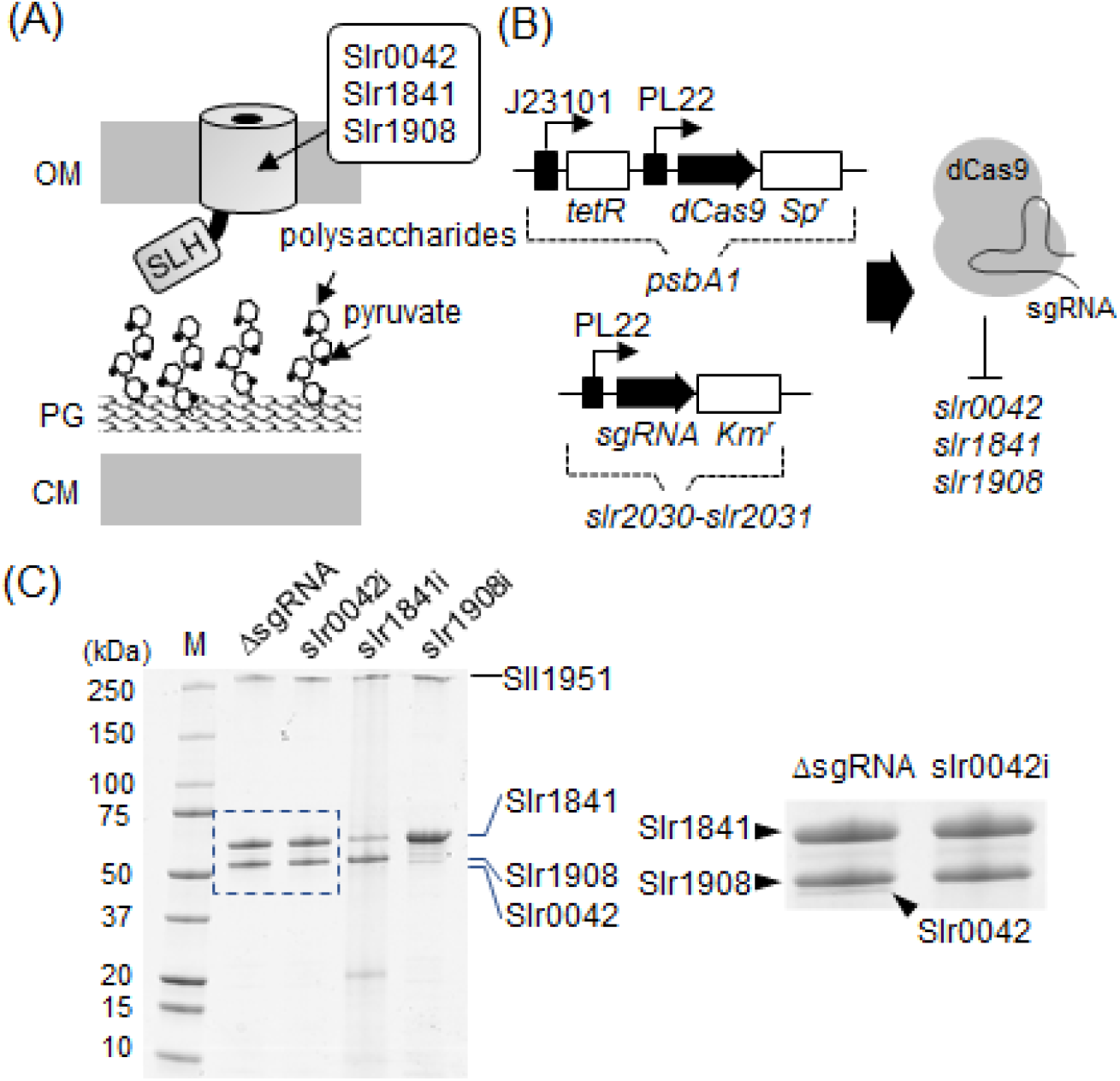
Downregulation of outer membrane proteins Slr0042, Slr1841, and Slr1908 proteins. (A) Schematic representation of PCC 6803 cell surface structure. Slr0042, Slr1841, and Slr1908, which represent approximately 80% of the outer membrane proteins, bind to peptidoglycan-linked polysaccharides via their periplasm-exposed SLH domains. Pyruvates are covalently bound to the polysaccharides. (B) The CRISPRi system. The dCas9 cassette, which comprises *tetR, dCas9*, and the spectinomycin resistance gene (*Spr*) were inserted at the *psbA1* neutral site of PCC 6803 chromosomal DNA. Transcription of the *tetR* and *dCas9* genes was controlled by the constitutive J23101 promoter and the aTc-inducible PL22 promoter, respectively. The sgRNA cassette, which comprises *sgRNA* and a kanamycin-resistance gene (*Kmr*), was inserted at the neutral site of *slr2030-slr2031*. The dCas9 protein and target-specific sgRNA form a complex that inhibits the expression of the target gene. (C) SDS-PAGE analysis of outer membrane proteins. Proteins (5 μg each) were analyzed using a 12.5% gel and stained with Coomassie Brilliant Blue (CBB). M, molecular mass standards. The boxed region of the gel was enlarged and is shown in panel on the right.

Here, we deprived the outer membrane of PCC 6803 by detaching it from the peptidoglycan, and characterized the effects on cell function. We demonstrated that outer membrane-deprived cells maintained photoautotrophic growth but liberated periplasmic and thylakoid luminal components into the external environment. These liberated substances sufficed to sustain the proliferation of heterotrophic bacteria in the absence of exogenous carbon sources. These findings show that breaching the outer membrane, without manipulating the cytoplasmic membrane, converted a free-living cyanobacterium into a chloroplast-like organism that conducts photosynthesis and released its biogenic materials to the extracellular space. This conversion represents one of the possible examples of metabolic exploitation of a cyanobacterial endosymbiont, and provides a potential explanation of why the outer membrane markedly changed during the early stages of chloroplast evolution. Further, this quasi-chloroplast provides the opportunity to exploit cyanobacterial photosynthesis to produce, for example, feedstock for biomanufacturing processes that have harnessed the heterotrophic bacteria.

## Results

### A Slr1841-repressed mutant exhibits outer membrane detachment while maintaining photoautotrophic growth

Approximately 80% of the outer membrane proteins of PCC 6803 comprise Slr1841 and its paralogs Slr1908 and Slr0042 (Fig. 1) (12). Reducing the amount of these proteins is assumed to destabilize or deprive the outer membrane because of loss of linkage to the peptidoglycan. We previously found that disruption of *slr1841* or *slr1908* genes led to induction of their cryptic paralogous genes (e.g., *sll1550*) and other outer membrane proteins (12). To avoid this undesirable inductions, we constructed mutant strains in which the expression of *slr1841, slr1908*, or *slr0042* was conditionally repressed (designated strains slr1841i, slr1908i, and slr0042, respectively). For this purpose, we employed a CRISPR-interference (CRISPRi) system (21) in which nuclease-deficient Cas9 (dCas9) and single guide RNA (sgRNA) form a complex that binds to a target gene, thereby inhibiting its transcription. Expression of dCas9 and sgRNA was controlled by the anhydrotetracycline (aTc)-inducible P_L22_ promoter (Fig. 1B). SDS-PAGE analysis revealed that in this system, the levels of Slr1841, Slr1908, and Slr0042 proteins were reduced by >50%, compared with the control strain whose chromosomal DNA harbors the same genetic construct without sgRNA (ΔsgRNA) (Fig. 1C). Note that the amount of Slr1841 protein in slr1908i strain unexpectedly increased by ca. 1.9-fold via an unknown mechanism. The ratio of total abundance of Slr1841, Slr1908, and Slr0042 proteins in the ΔsgRNA, slr0042i, slr1841i, and slr1908i strains was approximately 1:0.95:0.54:1.18, respectively. The extent of repression of Slr1841, Slr1908, or Slr0042 proteins was stable during culture days two to seven (Fig. S1).

As expected, the outer membrane of slr1841i strain was detached from the peptidoglycan (Fig. 2AB). Electron microscopy of thin-sectioned cells revealed that 72.5 ± 18.3% of cell surface was deprived of the outer membrane (n = 20, total examined cell surfaces in lengths = 82.4 μm). Strains slr1908i and slr0042i did not exhibit detectable alterations of cell surface structure. The photoautotrophic proliferation was maintained in slr1841i strain, although the specific growth rate was modestly reduced to 0.72 day^-1^ (during days 1 to 3) compared with ~1.10 day^-1^ for ΔsgRNA, slr0042i, and slr1908i strains (Fig. 2C). When slr1841i strain was transferred to a fresh medium containing aTc after three days of culture, the specific growth rate kept similar value (0.71 day^-1^), and this value did not significantly change during days 0 to 3 (data not shown).

**Fig. 2.**
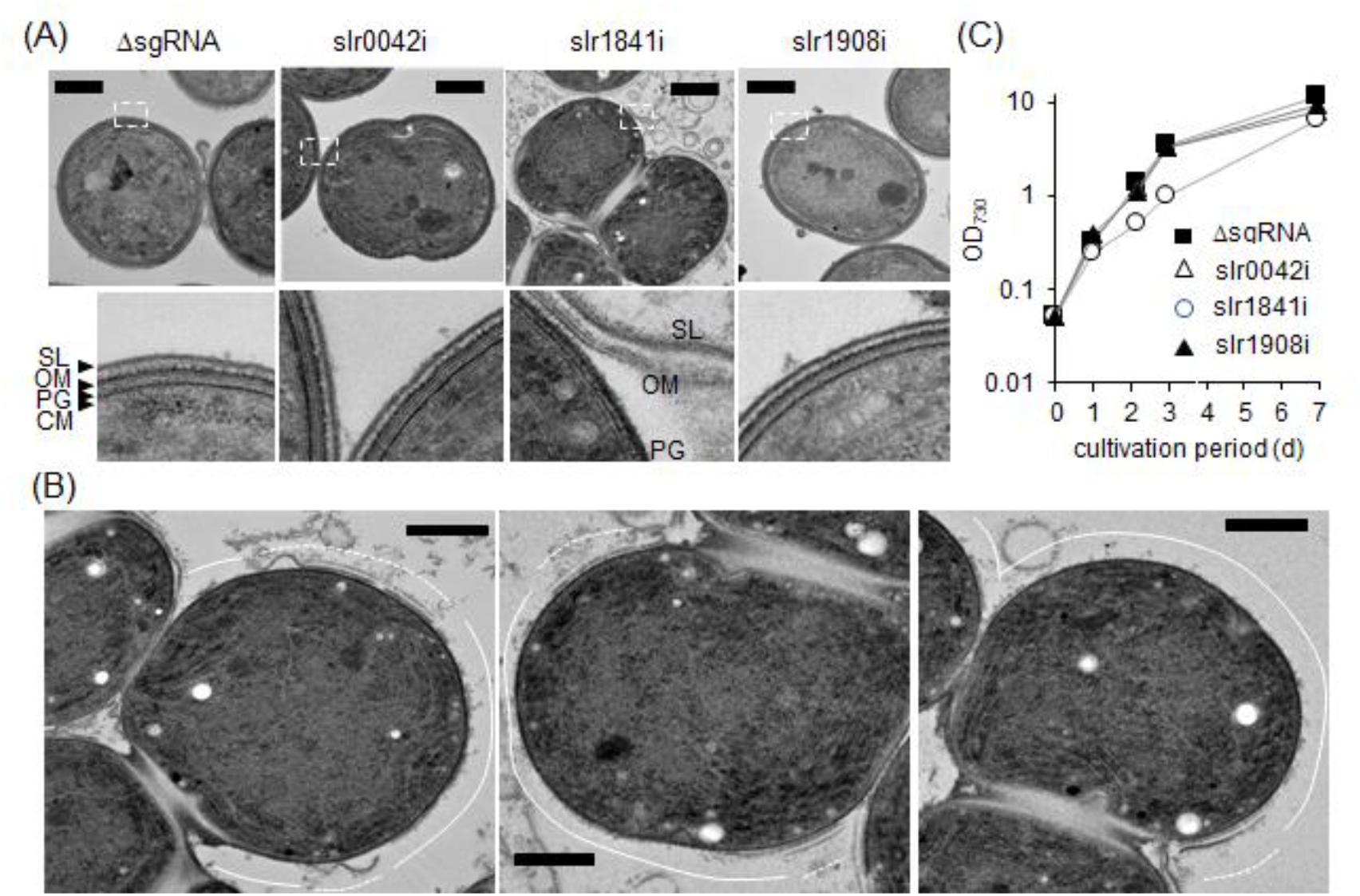
Cell surface structures and growth curves of ΔsgRNA, slr0042i, slr1841i, and slr1908i strains. (A and B) Electron micrographs of ultra-thin sectioned cells stained with 2% uranyl acetate and 0.4% lead citrate. Boxed regions are enlarged in the lower panels. Bars = 500 nm. Abbreviations: SL, S-layer; OM, outer membrane; PG, peptidoglycan; CM, cytoplasmic membrane. Three representative micrographs of strain slr1841i are enlarged and shown in (B). Solid and dashed lines indicate the regions where the outer membrane was detached or remained attached to the cell, respectively. Bars = 200 nm. (C) Growth curves of ΔsgRNA, slr0042i, slr1841i, and slr1908i strains. Data represent the mean of three independent experiments.

To determine whether outer membrane deprivation affected the photosynthetic activity, we employed pulse-amplitude modulation (PAM) measurement of chlorophyll fluorescence (Table S3 and Fig. S2) (22). The overall trace of chlorophyll fluorescence of slr1841i and ΔsgRNA strains were similar. There was no significant difference in the Fv/Fm value, which represents the maximum quantum yield of photosystem (PS) II, suggesting that the photosynthetic apparatus of slr1841i strain remained mostly intact. We note that, however, continuous light illumination modestly decreased the quantum yield of PSII (Fv’/Fm’ value) of this strain; hence, possible disturbance of photosynthetic electron flow could not be ruled out. This notion is discussed later in this paper.

### Weakening the peptidoglycan–SLH domain interaction leads to a slr1841i-like phenotype

To confirm that outer membrane detachment was caused by the reduction of the physical linkage to the peptidoglycan and not that of the Slr1841-specific function, we weakened the interaction between SLH domains of slr0042, slr1841, and slr1908 with the peptidoglycan. Mesnage et al. (23) reported that in *Bacillus anthracis*, pyruvylation of the peptidoglycan-linked polysaccharides, for which *csaB* is responsible, is required for binding with the SLH domain. The authors further presumed that the same mechanism applies to the SLH domain-peptidoglycan interaction in cyanobacteria. Accordingly, we first examined the occurrence of the pyruvylation of peptidoglycan-linked polysaccharides, and we detected ca. 6 μg/mg pyruvate in the peptidoglycan preparation (Fig. 3A). Next, we used the CRISPRi system to downregulate the expression of *slr0688*, which is the only gene homologous to *csaB* in the PCC 6803 genome. While no detectable change in the composition of outer membrane proteins was observed, the amount of pyruvate was decreased by approximately twofold in the *slr0688*-repressed strain (slr0688i) (Fig. 3A, B). To characterize the interaction between a pyruvate-depleted peptidoglycan and the SLH domain, we prepared recombinant SLH domains of Slr0042, Slr1841, and Slr1908, which were fused to glutathione *S*-transferase (GST-SLH). The purified GST-SLH preparations were mixed with purified peptidoglycan preparations, and the proteins that remained in the supernatant were analyzed after removing peptidoglycan-bound GST-SLHs (Fig. 3C). As judged by protein band intensity on SDS-PAGE gel, GST-SLHs were removed from the supernatant by ca. 80% when mixed with the peptidoglycan of ΔsgRNA. In contrast, it was removed only about 30% when mixed with the peptidoglycan of slr0688i strain, showing the lower-affinity interactions between GST-SLHs and the pyruvate-reduced peptidoglycan. Electron micrographs of thin-sectioned slr0688i cells showed detachment of the outer membrane, similar to that of slr1841i strain (Fig. 3D). Further, the growth rate and the chlorophyll fluorescence profiles were similar to those of slr1841i strain (Fig. 3E and Fig. S2). Accordingly, we concluded that outer membrane deprivation was caused by the impairment of the physical linkage between SLH domains and the peptidoglycan.

**Fig. 3.**
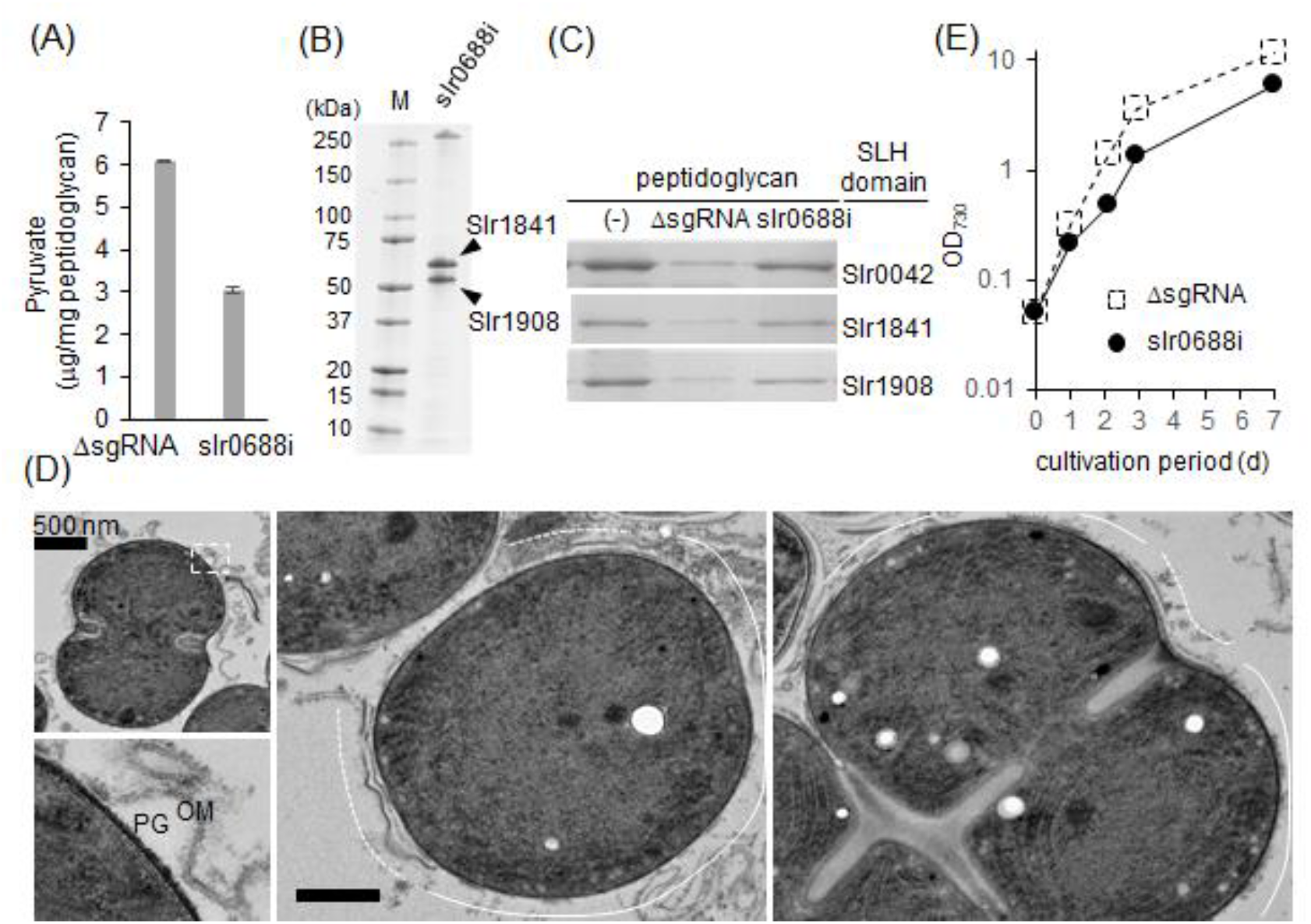
Phenotype of slr0688i strain. (A) Quantitation of pyruvate covalently bound to the peptidoglycan-linked polysaccharides in ΔsgRNA and slr0688i strains. Data represent the mean of three independent experiments ± SD. (B) SDS-PAGE analysis of outer membrane proteins. Proteins (5 μg each) were analyzed using a 12.5% gel and stained with using CBB. M, molecular mass standards. (C) Peptidoglycan-binding assay of GST-SLHs. Purified GST-SLH proteins were mixed with purified peptidoglycans of strains ΔsgRNA and slr0688i. Proteins remained in the supernatant were collected by centrifugation at 20,000 × *g* for 20 min at room temperature and analyzed using SDS-PAGE. Gels were stained with CBB. (D) Electron micrographs of ultra-thin sectioned slr0688i cells stained with 2% uranyl acetate and 0.4% lead citrate. Boxed regions are enlarged in the lower panel. Bars = 500 nm. Abbreviations: OM, outer membrane; PG, peptidoglycan. Two representative micrographs are enlarged and shown in the right panels. Solid and dashed lines indicate the regions where the outer membrane is detached or remains attached to the cell, respectively. Bars = 200 nm. (E) Growth curve of strain slr0688i. Data represent the mean of three independent experiments. The growth curve of strain ΔsgRNA is taken from Fig. 2C (dashed line).

### Outer membrane-deprived cells liberate periplasmic and thylakoid luminal components

The outer membrane deprivation was assumed to cause the release of cellular materials, such as periplasmic components. We first analyzed the proteinaceous components in culture supernatants of slr1841i and slr0688i strain (Fig. 4). As expected, the amount of proteins in slr1841i and slr0688i supernatants increased up to ~0.35 g/L during five days of culture (Fig. 4A). Wide variety of proteins was present in these supernatants (Fig. 4B). Mass-spectrometric analysis identified 82 and 100 proteins in slr1841i and slr0688i supernatants, respectively, among which 78 were identified in both supernatants (Tables 1, S4, S5). Signal peptides were detected in 83% and 71% proteins from slr1841i and slr0688i strains, respectively. These proteins contained periplasmic marker proteins such as *N*-acetylmuramoyl-L-alanine amidase (24), and of 57 periplasmic proteins identified in the study by Fulda et al. (24), 28 and 27 proteins were present in the slr1841i and slr0688i supernatants, respectively. These findings suggest that a large portion of the proteins released from these strains consists of the periplasmic components. The reason for presence of the proteins lacking a signal peptide is currently unknown, but cell lysis of slr1841i and slr0688i strains seems unlikely because the abundant cytoplasmic proteins, e.g., ribulose-1,5-bisphosphate carboxylase/oxygenase or other cellular metabolic enzymes, were absent in their culture supernatants. Notably, we identified plastocyanin, photosystem II 12 kDa extrinsic protein (psbU or Sll1194), photosystem II manganese-stabilizing polypeptide (psbO or Sll0427), and Slr0924 protein (25, 26), each of which functions in the thylakoid lumen. Given the absence of cell lysis, this finding suggests some degree of continuity throughout the thylakoid lumen-periplasm-extracellular environment, although connectivity between thylakoid and cytoplasmic membranes is the subject of longterm debate (27–29).

**Fig. 4.**
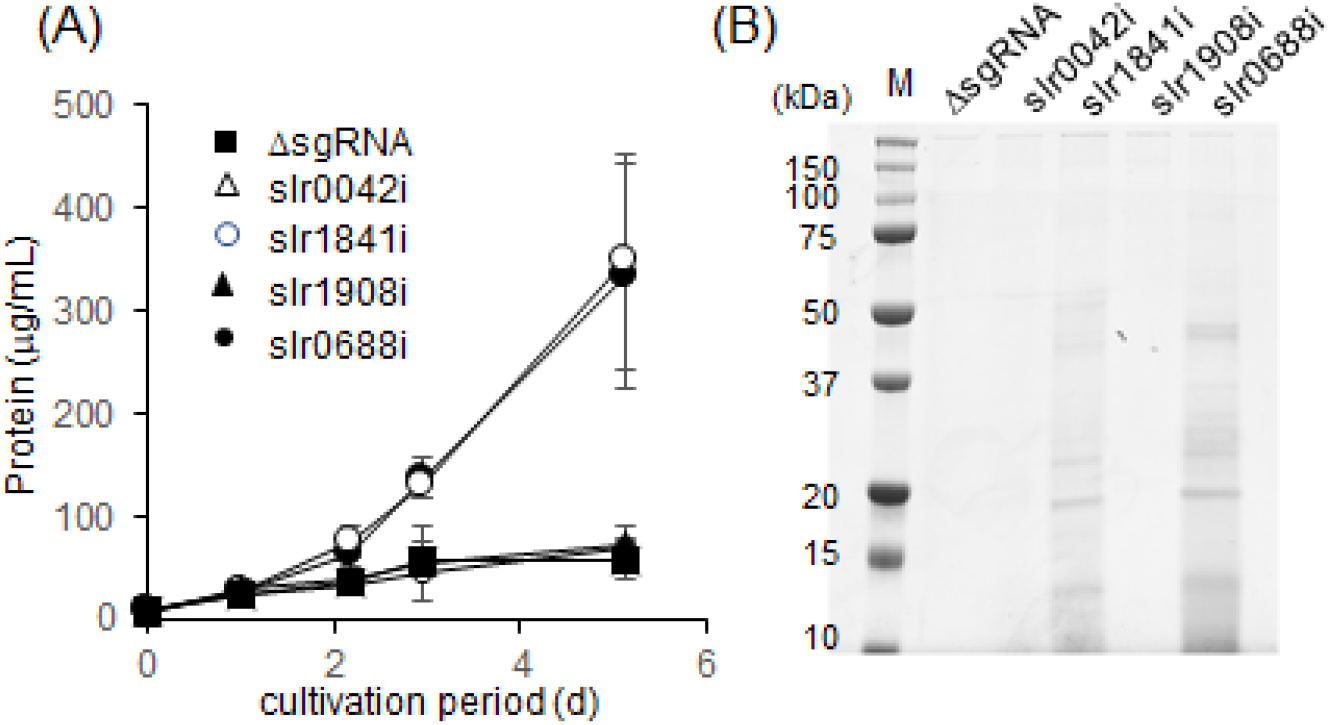
Proteins in culture supernatants of ΔsgRNA, slr0042i, slr1841i, slr1908i, and slr0688i strains. (A) Protein concentrations of supernatants. Data represent the mean of three independent experiments ± SD. (B) Proteins in 5 days-cultured supernatants. Twenty microliter of the supernatants were analyzed by SDS-PAGE, stained with CBB. M, molecular mass standards

**Table 1.** Proteins detected in culture supernatants of slr1841i and slr0688i strains

| Strain | Identified proteins | Proteins with signal peptide<sup>a</sup> | Proteins identified in both supernatants |
| --- | --- | --- | --- |
| slr1841i | 82 | 83% | 78 |
| slr0688i | 100 | 71% |  |

<sup>a</sup> Presence of signal peptide was analyzed using SignalP 5.0

### The photosynthetic electron transport chain is accessible to external hydrophilic oxidizing agent in outer membrane-deprived cells

To test whether the thylakoid lumen of slr1841i and slr0688i strains was accessible to the extracellular space, we determined if an exogenously added hydrophilic oxidizing agent was reduced by the photosynthetic electron transport chain, which resides on the thylakoid membrane. We therefore exposed cells to such a compound, 0.1 mM KMnO_4_, which becomes decolorized when reduced (Fig. 5). When the slr1841i or slr0688i-KMnO_4_ suspension was illuminated, a rapid decolorization was observed. In contrast, only a modest decolorization occurred in the dark, suggesting that KMnO_4_ was mainly reduced by photosynthetically generated electrons but not by other cellular constituents. The decolorization of the ΔsgRNA-KMnO_4_ suspension was much slower than that of slr1841i or slr0688i-KMnO_4_ suspension either in illuminated or dark conditions. These observations suggest that slr1841i and slr0688i strains allow the movement of KMnO_4_ between their thylakoid lumen and extracellular space. We therefore concluded that in slr1841i and slr0688i strains, the thylakoid lumen and the extracellular space are spatially connected, although the connection may be transient.

**Fig. 5.**
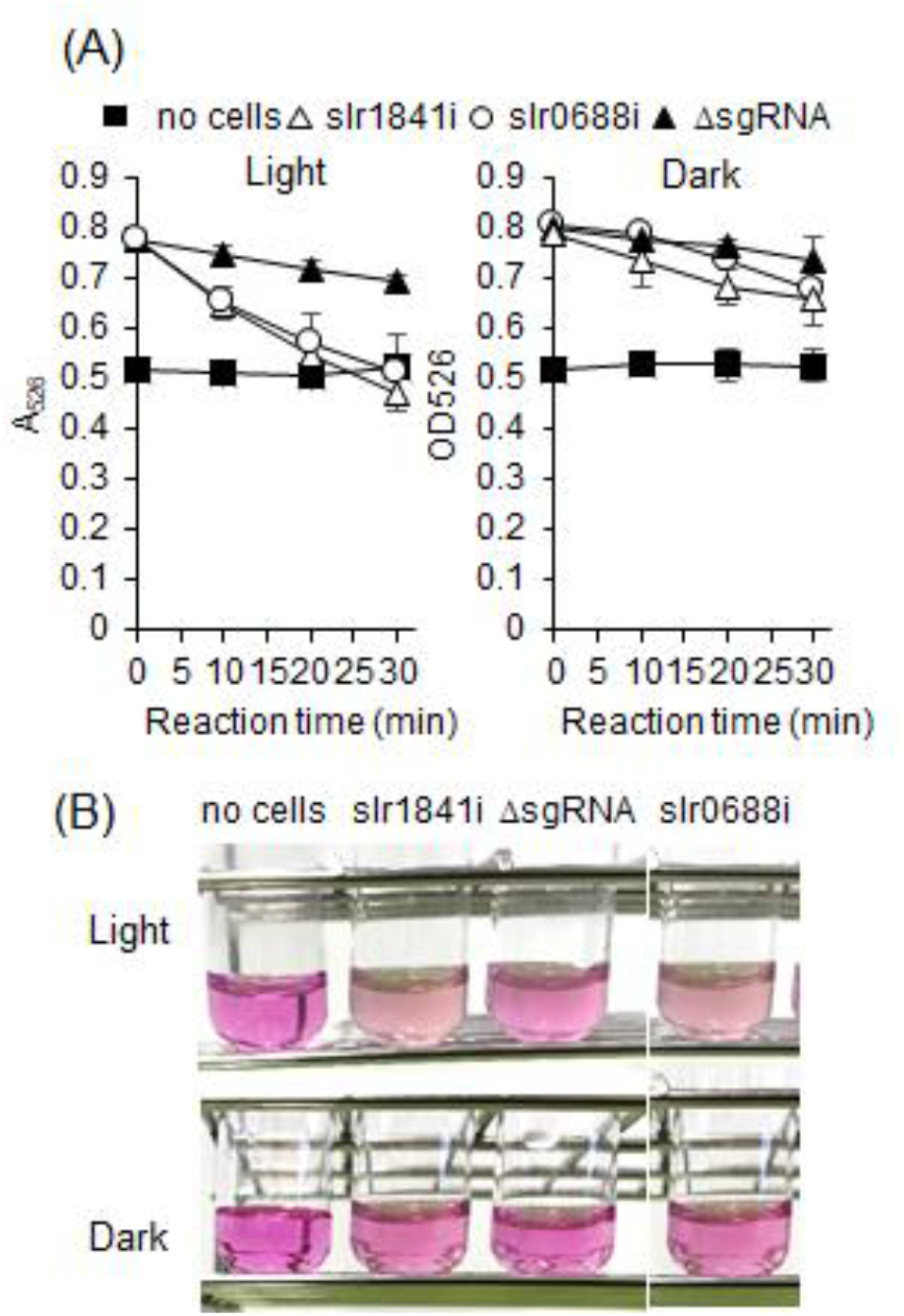
Reduction of KMnO_4_ by the photosynthetic electron transport chain. (A) Reduction of KMnO_4_ by ΔsgRNA, slr1841i, and slr0688i strains. Cells cultured for 3 days were suspended in 50 mM K-P buffer, pH 7.4, and mixed with 0.1 mM KMnO_4_. The mixture was incubated at 30 °C with shaking, with or without light illumination (100 μmol/m^2^/s). Reduction of KMnO_4_ was monitored by measuring A_526_. Data represent the mean of three independent experiments ± SD. (B) Representative images of reaction mixtures incubated for 30 min.

### Metabolomic profile of the cell culture supernatant

Metabolites in the culture supernatant of ΔsgRNA, slr1841i, and slr0688i strains were analyzed using capillary electrophoresis-time-of-flight mass spectrometry (CE-TOF/MS). Of the 43 identified compounds (Table S6), supernatants of ΔsgRNA, slr1841i,and slr0688i strains contained 34, 40, and 35 compounds, respectively. The abundant cytosolic compounds, such as ATP or NADP(H), were not detected. This suggests the absence of cell lysis, consistent with the results of proteomic analysis described above. Of above compounds, 11 were present with more than three-fold higher concentration in the supernatants of slr1841i and slr0688i strains compared to that of ΔsgRNA strain, whereas three compounds were unique to the supernatant of strain ΔsgRNA (Table 2). Notably, of the 11 compounds, five are nucleotide-related compounds such as adenosine and guanosine. This was consistent with the previous report showing that the thylakoid lumen contains nucleotide-related metabolic activities (30–32). The metabolites detected in the supernatants of slr1841i and slr0688i strains may be linked to the periplasmic and thylakoid luminal activities.

**Table 2.** Comparison of metabolites detected in strains ΔsgRNA, slr1841i, and slr0688i

| Metabolite<sup>a</sup> | Concentration (μM) | | |
| --- | --- | --- | --- |
| | ΔsgrRNA | slr1841i | slr0688i |
| 2'-Deoxycytidine | ND | 1.1 | 1.3 |
| 3-Hydroxybutyric acid | ND | 1.3 | 12 |
| 5-Oxoproline | ND | 0.7 | 1.4 |
| Acetoacetic acid | ND | 1.5 | 1.5 |
| Adenosine | ND | 0.8 | 0.6 |
| Cytidine | 0.1 | 0.7 | 0.5 |
| Glutamic acid | ND | 0.3 | 0.2 |
| Guanosine | 0.2 | 1.5 | 0.8 |
| *N*<sup>6</sup>-Methyl-2'-deoxyadenosine | ND | 0.4 | 0.3 |
| *p*-Aminobenzoic acid | ND | 1.1 | 1.1 |
| Phenylalanine | ND | 0.2 | ND |
| 2-Oxoglutaric acid | 3.9 | ND | ND |
| Glycolic acid | 22 | ND | ND |
| Nicotinic acid | 0.3 | ND | ND |

ND, not detected
<sup>a</sup> Metabolites present with more than three-fold higher or lower concentration in the supernatants of slr1841i and slr0688i strains compared to that of $\Delta$ sgrRNA strain.

### Liberated materials from outer membrane-deprived cells serve as nutrients for heterotrophic bacteria

On the basis of the calculation using the general protein carbon content of 0.52 g/g protein (33), the liberated materials (ca. 0.35 g proteins per liter) from the five days-cultured outer membrane-deprived cells were estimated to contain at least 0.18 g/L of organic carbons, which we assumed was sufficient to sustain the growth of heterotrophic bacteria without adding exogenous carbon sources. To test this possibility, we inoculated *E. coli, B. cereus, Pseudomonas aeruginosa*, and *Staphylococcus aureus* to five days-cultured supernatants of ΔsgRNA, slr1841i, and slr0688i strains, and incubated at 30°C with shaking (Fig. 6). These bacteria proliferated in the supernatants of slr1841i and slr0688i strains, but no obvious growth was observed in the supernatant ofΔsgRNA strain. This indicates that biogenic materials released from the outer membrane-deprived cells served as organic nutrients for these heterotrophic organisms.

**Fig. 6.**
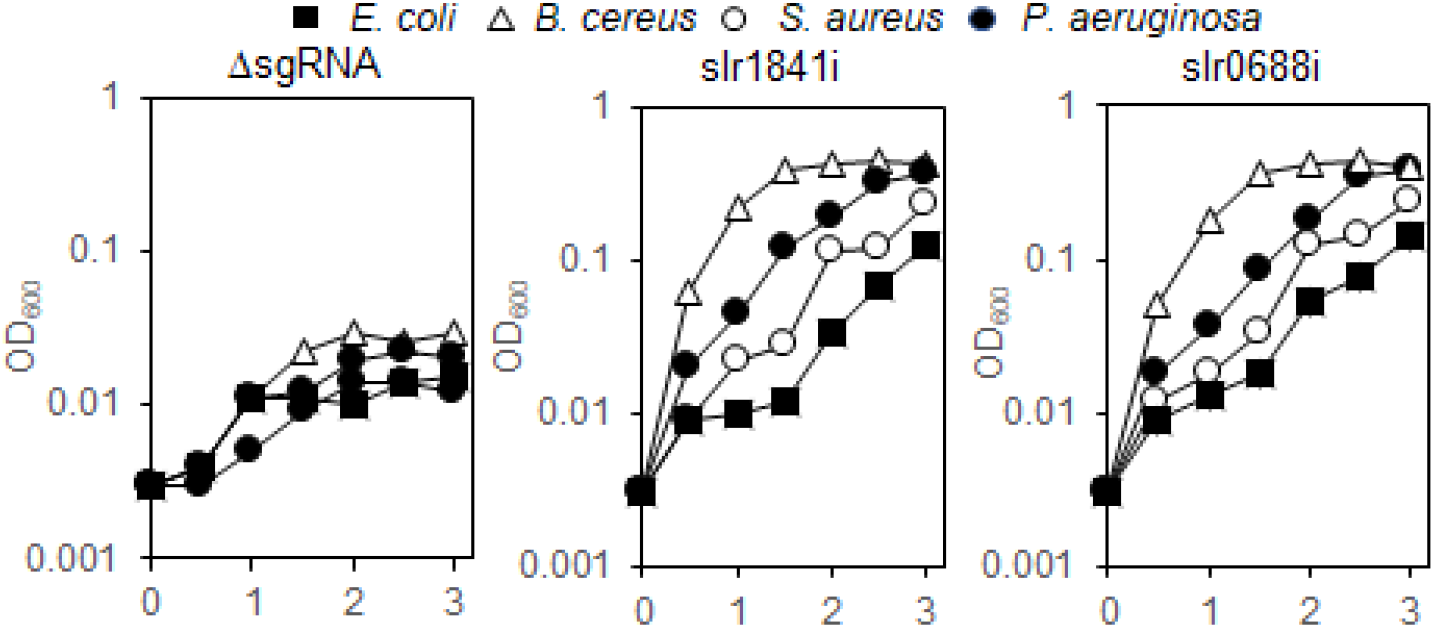
Growth of heterotrophic bacteria in culture supernatants of ΔsgRNA, slr1841i, and slr0688i strains. *E. coli, B. cereus, S. aureus*, and *P. aeruginosa* were inoculated to supernatants of 5 days-cultures of ΔsgRNA, slr1841i, and slr0688i strains, and the growth was monitored by measuring OD600. Data represent the mean of three independent experiments.

## Discussion

It has been considered that the cytoplasmic membrane of cyanobacteria is the primary barrier that must be overcome to exploit the photosynthesis for establishing the endosymbiotic relationship with the heterotrophic host cell (4, 34). The present study, on the other hand, we demonstrated that depriving the cyanobacterial outer membrane, without altering the cytoplasmic membrane, causes the release of cellular organic constituents that sufficed to sustain the proliferation of heterotrophic bacteria in the absence of exogenous carbon sources.

The interaction between the outer membrane and the peptidoglycan of Gram-negative bacteria has mainly been studied in the phylum Proteobacteria using *Escherichia coli* or *Salmonella typhimurium* as a model (19)]. These bacteria utilize multiple factors for forming the outer membrane-peptidoglycan linkage as follows: (i) Murein-lipoprotein Lpp, covalently linked to the peptidoglycan, anchors the outer membrane via its lipid moiety (15, 16); (ii) The highly abundant outer membrane protein OmpA binds to the peptidoglycan via its C-terminal periplasmic domain (35, 36); and (iii) The trans-envelope complex Tol-Pal, which comprises (at least) TolQ TolA, TolR, TolB, and Pal, contributes to the proper invagination of the outer membrane (37–40). In contrast, these factors are not conserved among the Cyanobacteria; to date the SLH domain–peptidoglycan interaction is the only known mechanism (11, 12). Deletion of *lpp* or *tol-pal* of *E. coli* destabilizes the outer membrane, leading to the formation of blebs and leakage of periplasmic enzymes (15, 38, 41). However, the entire outer membrane remains loosely associated with the peptidoglycan and surrounds the cell surface. Under normal growth conditions, mutants with multiple deletions in genes encoding above factors cannot be isolated (42), indicating that the outer membrane is essential for proper cell growth. This is in clear contrast to the case of PCC 6803, in which impairment of the SLH domain–peptidoglycan interaction leads to >70% removal of the outer membrane, while the cell growth was not severely inhibited (Fig. 2, 3). A possible explanation of the requirement of the outer membrane for the growth of *E. coli* is the involvement of outer membrane proteins in controlling the biosynthesis and hydrolysis of peptidoglycan, e.g., LpoA and LpoB regulate the activities of the peptidoglycan synthases PBP1a and PBP1b (43), and NlpI regulates peptidoglycan hydrolases (44). Extensive detachment of the outer membrane may alter the regulation of peptidoglycan biosynthesis, hydrolysis, or both, mediated by LpoA, LpoB, and NlpI, possibly leading to impaired cellular integrity. In contrast, neither *lpoA* or *lpoB* is conserved in cyanobacteria (protein sequences > 30% identity, > 50% sequence coverage, E-value < e-5 was considered as homologs), and the NlpI-homolog PratA does not localize to the outer membrane (45). Hence, the effect of detachment of the outer membrane on peptidoglycan biosynthesis/hydrolysis may be less severe compared with that of *E. coli*. Similar observations were made with bacteria belonging to the class *Negativicutes* that lack *lpp, ompA, tol-pal, lpoA, lpoB*, and *nlp* but employ SLH domain–peptidoglycan interactions resembling those of cyanobacterial species; weakening of this interaction leads to outer membrane detachment without significantly inhibiting cell growth (14, 46). The mechanism that controls peptidoglycan metabolism in cyanobacteria and *Negativicutes* may therefore differ from that of Proteobacteria, and it requires further research.

An unexpected finding of the present study was that the proteins liberated from slr1841i and slr0688i strains contained thylakoid luminal components. The thylakoid membrane closely contacts the cytoplasmic membrane at a defined region called the convergence zone (or thylakoid center), where multiple thylakoids converge (27, 28). This region is closely tethered through an unknown mechanism to the cytoplasmic membrane and serves as a site of biogenesis of thylakoid proteins, coupled with the incorporation of cofactors from the cytoplasmic membrane and the periplasm. Given that the cytoplasmic membranes of slr1841i and slr0688i strains remain mostly intact as judged by the absence of cell lysis, the convergence zone seems a probable site through which thylakoid luminal components exit and exogenous KMnO_4_ enters. Although the latest cryo-electron tomography study did not provide evidence of the physical fusion of the thylakoid lumen and the periplasm (27), our present results suggest that the convergence zone allows, at least transiently, the diffusional movement of certain cellular molecules between them. Leakage of thylakoid luminal components, particularly plastocyanin, which is a component of the photosynthetic electron transport chain, may decrease the efficiency of transport of photosynthetically generated electrons (22). This notion is consistent with findings that the Fv’/Fm’ values of strains slr1841i and slr0688i were modestly lower than that of strain ΔsgRNA.

Breaching the outer membrane barrier can be achieved by weakening the SLH domain–peptidoglycan interaction as well as by other changes that permeabilize the outer membrane, such as introducing channel proteins with a large pore size. Thus, CppS/F proteins of the muroplast seem good candidates for serving this purpose. Further, considering the extensive changes of cyanobacterial outer membrane during the generation of the primitive chloroplast, the barrier function of the cyanobacterial outer membrane appears likely to have been breached during the earliest stage of the chloroplast evolution, and this events may represent a possible example of exploiting the cyanobacterial endosymbiont. In this regard, it is of interest to analyze the outer membrane of the *Paulinella chromatophora* chloroplast, which retains peptidoglycan and the outer membrane and is therefore referred to as “ongoing” chloroplast generation, although this organism is phylogenetically unrelated to Archaeplastida (47).

Lastly, we propose that research on the cyanobacterial outer membrane not only will shed light on the early stage of the evolution of chloroplast generation, but also provide potential opportunities to exploit cyanobacterial photosynthesis for industrial applications. The liberated biogenic materials from outer membrane-deprived cyanobacterial species can be obtained without harvesting and disrupting the cells and are also usable for feedstock for biomanufacturing using heterotrophic bacteria. Although the amount of liberated materials was not currently high enough to support the rapid growth of heterotrophic bacteria, the application of ever growing sets of molecular-genetic tools can aid for further improving the productivity to enhance the potential of this quasi-chloroplast.

## Materials and Methods

### Strains and culture conditions

PCC 6803 and its mutants were cultured in BG11 medium buffered with 4.5 g/L TES (pH 7.5), at 30 °C with shaking, under continuous light illumination (100 μmol/m^2^/s). The CRISPRi system was induced by adding 1 μg/mL aTc. *E. coli* K-12 (strain 4401 from Coli Genetic Stock Center, Yale University), *B. cereus* (ATCC 14579), *P. aeruginosa* (ATCC 10145), and *S. aureus* (ATCC 12600) were cultured in LB medium (10% tryptone, 5% yeast extract, 5% NaCl) supplemented with 5 mM MgCl2 at 30 °C and shaking, before using for the growth analysis in the culture supernatants of PCC 6803.

### Construction of CRISPRi strains

The dCas9 gene cassette which comprises the *tetR* gene, aTc-inducible promoter P_L22_, the *dCas9* gene, and a spectinomycin resistance gene *(Spr)*, was amplified from the chromosomal DNA of strain PCC 6803 LY07, using the primer set shown in Table S7 (21). By this PCR reaction, the dCas9 cassette was amplified with its downstream and upstream regions of the *psbA1* neutral site. PCR products (approximately 10 ng) were mixed with 300 μL of a PCC 6803 culture harvested at OD_730_ ~0.3, and transformants were selected on BG11 agar plates containing 25 μg/mL spectinomycin. Next, the sgRNA cassette, which comprises the P_L22_ promoter, sgRNA, and a kanamycin resistant gene (*Kmr*), was similarly amplified with its upstream region of slr2030 and downstream region of slr2031, from strain LY07. The *slr0042, slr1841, slr1908*, and *slr0688*-targeting sgRNA sequences shown in Table S8 were inserted into the sgRNA cassette using overlapping PCR. Potential off-target sites of these sgRNA sequences were analyzed using CasOT software (48). This analysis did not detect off-target sites with less than five sequence mismatches, indicating the potential off-target effect is insignificant. The PCR products were used to transform PCC 6803 cells harboring the dCas9 cassette as above. Finally, strains slr0042i, slr1841i, slr1908i, and slr0688i were selected and segregated on BG11 agar plates containing 25 μg/mL spectinomycin and 20 μg/mL kanamycin.

### Preparation of outer membranes

Cells were grown in BG11 medium containing 1 μg/mL aTc, and the outer membrane was isolated as previously described (12). ImageJ software was used to quantitate the intensity of protein bands on Coomassie Brilliant Blue (CBB)-stained SDS-PAGE gels.

### Electron microscopy

Cells were cultured for three days, frozen in liquid propane, and substituted with 2% glutaraldehyde, 1% tannic acid in ethanol, and 2% distilled water at −80 °C. After the samples were warmed to 4 °C, they were dehydrated in ethanol, soaked twice in propylene oxide for 30 min each, and added to a 70:30 mixture of propylene oxide and Quetol-651 resin (Nisshin EM, Tokyo, Japan) for 2 h. Samples were then added to 100% resin and polymerized at 60 °C for 48 h. The samples were thin-sectioned (70-nm thick), stained with 2% uranyl acetate and 0.4% lead citrate, and observed using a transmission electron microscope (JEM-1400Plus; JOEL Ltd, Tokyo, Japan) at an acceleration voltage of 100 kV

### Chlorophyll fluorescence measurement

Cells were cultured for three days, and then the cell density was adjusted to 1 μg/mL chlorophyll using BG11 medium. The cell suspension was then incubated in the dark for 15 min. Chlorophyll fluorescence was measured according to Misumi and Sonoike (49) using a WATER-PAM fluorometer (Waltz, Germany). Briefly, the minimum fluorescence level (Fo) was first determined at 650 nm. After a saturating light pulse (0.8 s) was delivered to confirm the photosynthetic activity, samples were illuminated with actinic light (660 nm, 562 μmol/m^2^/s), and a saturating light pulse was delivered 3–4 times to measure the maximum fluorescence intensity of the light-acclimated cells (Fm’). The maximum fluorescence level (Fm) was determined by adding 10 μM 3-(3,4-dichlorophenyl)-1,1-dimethylurea (DCMU) under illumination with actinic light. Fv and Fv’ are calculated as Fm-Fo, and Fm’-Fo, respectively.

### Quantification of pyruvylation of peptidoglycan-linked polysaccharides

Peptidoglycan-linked polysaccharides were prepared from cells cultured for three days using the method of Weckesser and Jurgens (50). Samples were hydrolyzed in 0.5 M HCl for 30 min at 100 °C, as described in the study Mesnage et al. (23). After adjusting the pH to 7.0 by adding NaOH solution, the amount of pyruvate was quantified using a Pyruvate Assay Kit (Sigma-Aldrich).

### Preparation of recombinant SLH domains and peptidoglycan-binding assay

The SLH domains of Slr0042 (amino acid residues 1–108), Slrl84l (residues 1–108), and Slr1908 (residues 1–126) fused with GST were expressed in *E. coli* BL21(DE3) and purified, as described in the study by Kojima et al. (11). The peptidoglycan-binding assay was performed as previously described (11).

### Sequence analysis

Sequence similarity searches were performed by BLAST program of the NCBI database. Putative signal peptides were predicted using SignalP-5.0 (http://www.cbs.dtu.dk/services/SignalP/).

### Proteomic analysis

Culture supernatants were collected from five days-cultures by centrifugation at 5,000 × *g* for 10 min at 4 °C, and were filtered through 0.45-μm-pore polyvinylidene difluoride (PVDF) membranes. Proteins were precipitated by adding ice-cold acetone (final concentration 80%) and incubated on ice for 4 h. Precipitates were collected by centrifugation at 20,000 × *g* for 20 min at 4 °C, washed once with 80% acetone, dried, and then solubilized in 100 mM Tris-HCl (pH 8.5) containing 0.5% sodium dodecanoate. Proteins (20 μg/20 μL solution) were reduced in 10 mM dithiothreitol for 30 min at 50 °C and alkylated by 30 mM iodoacetamide for 30 min at room temperature (RT). Cysteine (final concentration 60 mM) was added to terminate alkylation, and the solution was incubated for 10 min at RT. After adding 150 μL of 50 mM ammonium hydrogen carbonate, the proteins were digested using 400 ng of Lys-C and 400 ng of trypsin at 37 °C for overnight. Trifluoroacetic acid (5%) was added to the samples, and the supernatants were collected by centrifugation at 15,000 × *g* for 10 min at RT. Digested peptides were purified using a C18-spin column, dried, and solubilized in 3% acetonitrile-0.1% formic acid. The samples were analyzed using a liquid chromatography-mass spectrometry system comprising an UltiMate 3000 RSLCnano LC System (Thermo Fisher Scientific, MA) and a Q Exactive HF-X Hybrid Quadrupole-Orbitrap Mass Spectrometer (Thermo Fisher Scientific). In this system, the peptides were separated on CAPCELL CORE MP column (75 μm × 2.5 cm, particle size 2.7 μm, Osaka Soda, Okasa, Japan) using a step gradient of 0.1% formic acid (eluent A) and 0.1% formic acid in 80% acetonitrile (eluent B) at a flow rate of 200 nL/min. Timetable of the eluent B concentration was as follows: 0-4min, 2%; 4-27 min, 8%; 27-28 min, 44%; 28-34 min, 80%. MS scans were performed at 60,000 resolution with scan range of 495 to 865 *m/z*. Data was analyzed using Scaffold DIA Proteome software and the UniProtKB/Swiss-Prot database under following parameters: precursor tolerance, 8 ppm, fragment tolerance, 10 ppm, data acquisition type, overlapping DIA; peptide length, 8-70; maximum missed cleavages, 1; total peak intensity, >2.5E+8.0; false discovery rate, <1%.

### KMnO_4_ reduction assay

Three days-cultured cells were harvested by centrifugation at 5,000 × *g* for 10 min at RT, washed twice with 50 mM potassium phosphate buffer pH 7.4, suspended in the same buffer, and were adjusted to OD7_30_ = 0.1. KMnO_4_ was then added to a final concentration of 0.1 mM. This cell suspension was incubated at 30 °C with shaking with or without light illumination (100 μmol/m^2^/s), and reduction of KMnO_4_ was monitored by measuring the reduction of A_562_.

### Metabolomic analysis

Supernatants were collected from five days-cultures as described above, filtered through a 0.45-μm-pore PVDF membrane, and were filtered again through a 5-kDa cutoff centrifugal filter unit at 9,100 × *g* for 60 min. Samples were analyzed using Agilent CE-TOF/MS system (Agilent Technologies) with a fused silica capillary (50 μm × 80 cm) under following condition/parameters: CE voltage, 30 kV; MS ionization, electrospray ionization of positive or negative modes; MS scan range, 50-1,000 *m/z*. The data were analyzed using MasterHands software (ver.2.17.1.11) (Human Metabolome Technologies, Tsuruoka, Japan), using the Human Metabolome Technologies database to annotate the metabolites under following parameters: peak signal/noise ratio, >3; migration times tolerance, ± 0.5 min; *m/z* tolerance, ± 10 ppm. The concentration of the metabolites was determined on the basis of the relative peak areas compared with internal standard chemicals.

