## Supplementary material for "Outer membrane-deprived cyanobacteria liberate periplasmic and thylakoid luminal components that support the growth of heterotrophs": Figure S1 and S2

**SI TEXT**

### Figure legends

**Fig. S1.** Analysis of Slr1841 and Slr1908 in outer membrane preparations of ΔsgRNA, slr0042i, slr1841i, and slr1908i strains cultured for 2 and 7 days. Proteins (7 μg each) was analysed by SDS-PAGE, stained with CBB.

**Fig. S2** Chlorophyll fluorescence of ΔsgRNA, slr1841i and slr0688i strains.
Strains ΔsgRNA (A), slr1841i (B), and slr0688i (C) were analyzed using a WATER-PAM fluorometer. Measuring light (ML) and actinic light (AL) were turned on or off at the times indicated by arrowheads. DCMU was added as indicated by the arrows.

**SI Figures**

Figure S1

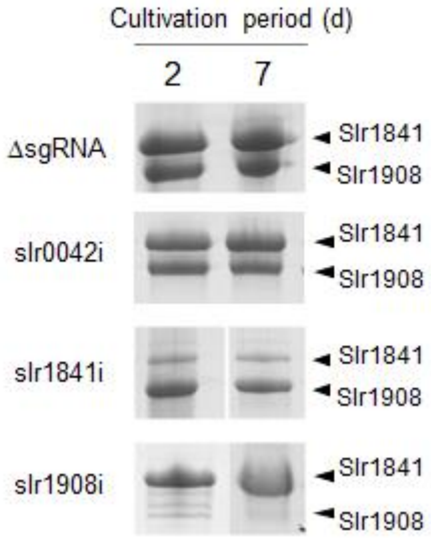

Figure S2

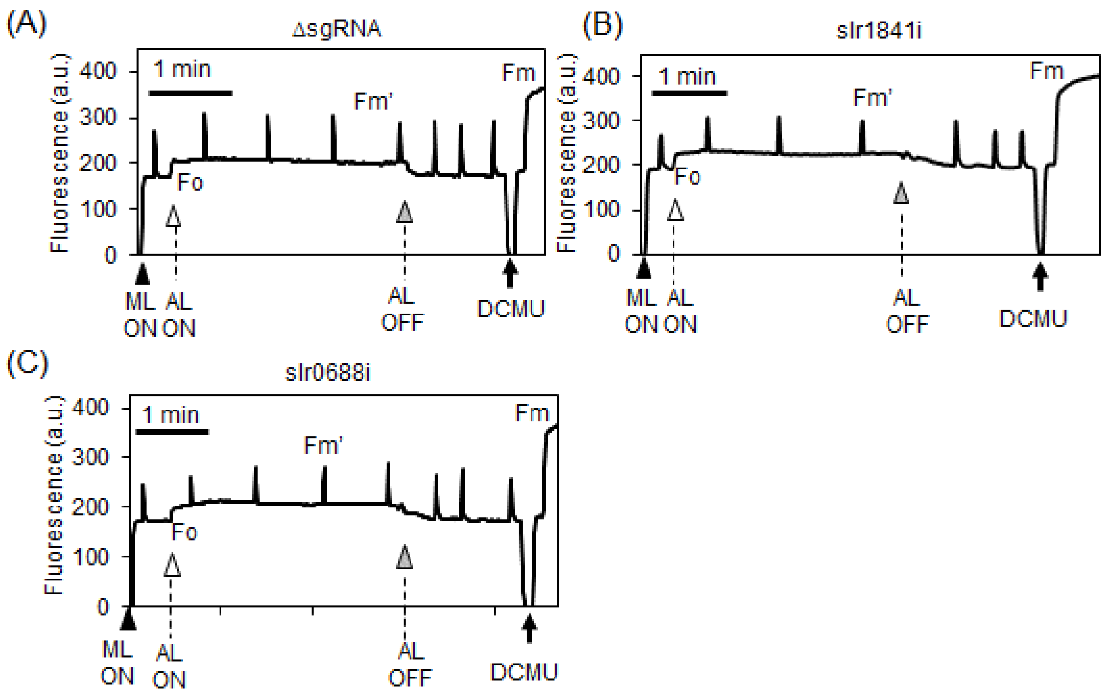

(A)
ΔsgRNA
Fluorescence (a.u.)
400
300
200
100
0
1 min
Fm'
Fm
Fo
ML ON
AL ON
AL OFF
DCMU
(B)
slr1841i
Fluorescence (a.u.)
400
300
200
100
0
1 min
Fm'
Fm
Fo
ML ON
AL ON
AL OFF
DCMU
(C)
slr0688i
Fluorescence (a.u.)
400
300
200
100
0
1 min
Fm'
Fm
Fo
ML ON
AL ON
AL OFF
DCMU
